# Connectome-wide search for functional connectivity locus associated with pathological rumination as a target for real-time fMRI neurofeedback intervention

**DOI:** 10.1101/2020.01.24.910430

**Authors:** Masaya Misaki, Aki Tsuchiyagaito, Obada A Zoubi, Martin Paulus, Jerzy Bodurka, Tulsa 1000 Investigators

**Author notes:** Corresponding authors Masaya Misaki, Jerzy Bodurka. The Tulsa 1000 Investigators include the following contributors: Robin Aupperle, Ph.D., Jerzy Bodurka, Ph.D., Justin Feinstein, Ph.D., Sahib S. Khalsa, M.D., Ph.D., Rayus Kuplicki, Ph.D., Martin P. Paulus, M.D., Jonathan Savitz, Ph.D., Jennifer Stewart, Ph.D., Teresa A. Victor, Ph.D.

## Abstract

Real-time fMRI neurofeedback (rtfMRI-nf) enables noninvasive targeted intervention in brain activation with high spatial specificity. To achieve this promise of rtfMRI-nf, we introduced and demonstrated a data-driven framework to design a rtfMRI-nf intervention through the discovery of precise target location associated with clinical symptoms and neurofeedback signal optimization. Specifically, we identified the functional connectivity locus associated with rumination symptoms, utilizing a connectome-wide search in resting-state fMRI data from a large cohort of mood and anxiety disorder individuals (N=223) and healthy controls (N=45). Then, we performed a rtfMRI simulation analysis to optimize the online functional connectivity neurofeedback signal for the identified functional connectivity. The connectome-wide search was performed in the medial prefrontal cortex and the posterior cingulate cortex/precuneus brain regions to identify the precise location of the functional connectivity associated with rumination severity as measured by the ruminative response style (RRS) scale. The analysis found that the functional connectivity between the loci in the precuneus (−6, −54, 48 mm in MNI) and the right temporo-parietal junction (RTPJ; 49, −49, 23 mm) was positively correlated with RRS scores (depressive, *p* < 0.001; brooding, *p* < 0.001; reflective, *p* = 0.002) in the mood and anxiety disorder group. We then performed a rtfMRI processing simulation to optimize the online computation of the precuneus-RTPJ connectivity. We determined that the two-point method without a control region was appropriate as a functional connectivity neurofeedback signal with less dependence on signal history and its accommodation of head motion. The present study offers a discovery framework for the precise location of functional connectivity targets for rtfMRI-nf intervention, which could help directly translate neuroimaging findings into clinical rtfMRI-nf interventions.

## 1. Introduction

Neurofeedback is a noninvasive technique of intervening in human brain activity (Sitaram et al., 2017). In neurofeedback training, a feedback signal of on-going brain activation is presented to participants to help them self-regulate their brain activity through learning to modulate the signal. A promising application of this intervention is a clinical treatment of psychiatric disorders via normalization of abnormal brain activation as a result of the training (Linhartová et al., 2019; Stoeckel et al., 2014; Young et al., 2017). Designing a neurofeedback protocol involves the identification of the target brain activation associated with a disorder symptom. In this regard, real-time fMRI neurofeedback (rtfMRI-nf) has a distinct advantage over other neurofeedback modalities, such as electroencephalography and/or near-infrared spectroscopy, in localizing a target region anywhere in the brain with high spatial resolution. fMRI is also the most popular neuroimaging method for mapping human brain function as well as altered function in a disease state. RtfMRI-nf can take advantage of the results of such fMRI functional mapping studies to determine the intervention target. The activation measure of rtfMRI-nf can either be activity at a specific region of interest (ROI), functional connectivity between regions, or a pattern of multiple regions’ activity (Watanabe et al., 2017). Thus, with its high spatial specificity and broad applicability of functional measurement, rtfMRI-nf can be a direct way of translating neuroscience knowledge into a clinical intervention.

In designing a rtfMRI-nf treatment protocol, abnormal brain activity associated with a disorder symptom is determined as a neurofeedback target, with the assumption that normalizing an abnormal brain activation could alleviate disorder symptoms. Two major approaches have been used for neurofeedback target determination (Sulzer et al., 2013). One approach involves referring to previous research characterizing disease-specific abnormal brain responses to identify a specific anatomical location associated with the disease. RtfMRI-nf can take advantage of the outcomes of abundant neuroimaging research, including systematic review and quantitative meta-analysis studies, to identify the neurofeedback target. This approach, however, cannot fully utilize the high spatial specificity of rtfMRI-nf. The result of a systematic review that summarizes studies with different specific aims based on a region name of atlas-based nomenclature does not indicate the exact location of abnormal brain activation. While quantitative meta-analysis, such as activation likelihood estimation (ALE) (Turkeltaub et al., 2002), can indicate a coordinate of the region involved in the disease, meta-analysis usually includes studies with broadly different aims or tasks. Thus, the identified locus may not be the region of specific functional abnormality but an overlap of blurred activation maps for different functions.

Another approach to identifying the rtfMRI-nf target is a functional localizer scan (Sulzer et al., 2013; Weiskopf et al., 2007). This approach performs a task that can elucidate a specific functional abnormality in fMRI and finds a locus of abnormal brain activation for each participant. This approach has a significant advantage in pinpointing the personalized target location, and researchers can fully take advantage of the high-spatial specificity of rtfMRI-nf. However, it is not always possible to fully utilize this approach since not all diseases have an established localizer task to identify abnormalities, and some tasks may not be applicable to patients with severe symptoms. For example, showing negative pictures repetitively to depressed patients could be harmful to their mood and possibly worsen their symptoms. Also, if the abnormality is expressed as a non-activation, we cannot locate the position of abnormality with a localizer scan (Young et al., 2018).

The present study introduced an alternative approach for data-driven and process-based neurofeedback target identification based on big data of resting-state fMRI, specifically for functional connectivity rtfMRI-nf. The data-driven approach can identify the exact location of abnormal brain activation by analyzing the original data, which is not performed in review or meta-analysis studies. Although the approach based on population data cannot personalize the target location like the functional localizer, it can identify the location with a non-active or low-connectivity abnormality through a comparison between disease and control groups. In addition, a resting-state fMRI scan is applicable to any patient population and could indicate an abnormality as a therapeutic target (Yamada et al., 2017).

Furthermore, if the dataset for identifying the target includes samples with a wide range of symptom spectrum measures across diagnostic groups, we can identify the locus of brain activation associated with symptom dimension. As the NIMH Research Domain Criteria (RDoC) framework highlighted (Morris and Cuthbert, 2012; Sanislow et al., 2010), current diagnostic systems for mental disorders are not based on neurobiological alterations, and the connection between diagnosis and underlaying neurobiology has not been established. The rtfMRI-nf intervention cannot be a full-fledged clinical treatment with such uncertainty and variability of neurobiological abnormality. A process-based framework of rtfMRI-nf (Lubianiker et al., 2019) has been proposed for precise intervention to accommodate such neurobiological variability in a diagnostic group. This framework identifies the neurofeedback target associated with a specific functional process of pathological abnormality instead of the average difference between disease and control groups. The data-driven approach can identify such a process-based target via direct access to the original data. A functional localizer cannot identify the locus associated with dimensional abnormality because the abnormality can be characterized only with a distribution of population data, not with one individual data point. Taken together, the data-driven process-based approach could be an optimal way to identify a neurofeedback target to make good use of the advantage of rtfMRI-nf.

To establish this approach, we need a dataset with large sample size, including a range of diagnostic groups and comprehensive measures of functional dimensions. The Tulsa 1000 study provides the ideal dataset for this purpose (Victor et al., 2018). The dataset includes both healthy participants and those with a psychiatric diagnosis with comprehensive measurements of biological and behavioral assessment, including fMRI and symptom scales. Using this dataset, the present study aimed to identify the functional connectivity locus associated with a specific symptom, rumination in mood and anxiety disorder participants, as a target of future rtfMRI-nf intervention.

Rumination has been defined as “the process of thinking perseveratively about one’s feelings and problems rather than in terms of the specific content of thoughts” (Nolen-Hoeksema et al., 2008). Rumination and associated repetitive negative thinking are a pervasive symptom observed in multiple psychiatric disorders, including depression, anxiety, substance abuse, obsessive-compulsive disorder, binge eating, and self-injurious behavior (McLaughlin et al., 2014; Wahl et al., 2019). Ruminative response style to a traumatic event also mediates the development of post-traumatic stress disorder symptoms (García et al., 2015). Rumination could exacerbate depression symptoms by enhancing the effect of depressed mood with repetitive thinking of negative thoughts and by interfering with problem solving (Nolen-Hoeksema et al., 2008). Ruminative response style is also a predictive factor of major depressive episodes (Spasojevic and Alloy, 2001). These indicate that rumination is a promising target for treatment to alleviate disorder symptoms as well as prevent symptom development across diagnoses. Indeed, cognitive behavioral therapies targeting rumination showed an effect of decreasing depressive symptoms (Jones et al., 2008; Schmaling et al., 2002; Watkins et al., 2011).

Neurobiologically, cortical midline structures (Nejad et al., 2013; Northoff and Bermpohl, 2004) involved in the default mode network (DMN) have been implicated in rumination and associated self-referential thinking (Burkhouse et al., 2017; Hamilton et al., 2015; Hamilton et al., 2011; Jiang et al., 2017; Johnson et al., 2009; Lois and Wessa, 2016; Murray et al., 2015; Nejad et al., 2013; Satyshur et al., 2018; Zhu et al., 2012; Zhu et al., 2017). Specifically, two core parts of the structure, the medial prefrontal cortex (MPFC) and the posterior cingulate cortex/precuneus (PCC/Prec) regions have been implicated in self-referential processing as well as pathological rumination due to their abnormal activity (Cooney et al., 2010; Nejad et al., 2019; Renner et al., 2015) and altered functional connectivity (Berman et al., 2014; Cheng et al., 2018; Connolly et al., 2013; Davey et al., 2017; Kuhn et al., 2012; Philippi et al., 2018; Yuan et al., 2018).

The current study searched for the precise locus of functional connectivity associated with rumination symptom severity for mood and anxiety disorder patients. The brain areas involved in this search included the MPFC and PCC/Prec regions. While studies indicate that functional connectivity in either the MPFC or PCC/Prec was associated with rumination, most of the studies used an *a priori*-defined seed region based on an anatomical atlas and did not perform a voxel-wise search for the seed and its connectivity. As these areas are not functionally homogeneous (Andrews-Hanna et al., 2010; Cavanna and Trimble, 2006; Leech et al., 2012), an *a priori* definition of the seed ROI could misidentify the precise location of the connectivity associated with rumination symptom. While Cheng et al. (2016) and Cheng et al. (2018) examined voxel-wise resting-state functional connectivity to find altered connectivity in participants with major depressive disorder compared to controls, they did not search for and determine connectivity correlated with rumination symptom severity. They, instead, searched the connectivity with group difference and evaluated a correlation with rumination in a post-hoc analysis. Thus, the precise locus of functional connectivity associated with rumination severity that can serve as a neurofeedback target with high-spatial specificity has not yet been identified.

To identify the precise location of the functional connectivity associated with rumination severity, the present study performed a connectome-wide association analysis (Misaki et al., 2018a; Shehzad et al., 2014). The connectome-wide analysis investigates comprehensive voxel-wise connectivity associations (Shehzad et al., 2014) utilizing multivariate distance matrix regression (MDMR) analysis (Anderson, 2001). This analysis examines voxel-wise connectivity association without *a priori* seed definition. We performed MDMR for resting-state fMRI data in the Tulsa 1000 study dataset with a regressor of rumination symptoms derived from the Ruminative Response Styles (RRS) scale (Nolen-Hoeksema and Morrow, 1991). We supposed that this analysis could enable us to identify the locus of functional connectivity significantly associated with rumination severity in mood and anxiety disorders, which can serve as a rtfMRI-nf target.

Additionally, we performed a simulation analysis to design an optimal real-time neurofeedback signal, following a framework introduced by Ramot and Gonzalez-Castillo (2018). This optimization with simulation is an additional benefit of accessing original data when designing a rtfMRI-nf treatment protocol. The simulation analysis was performed for the functional connectivity identified in the connectome-wide analysis described above. Two measures of online functional connectivity, sliding-window correlation and the two-point algorithm (Ramot et al., 2017), were evaluated. Through a data-driven search for the precise location of connectivity associated with rumination severity and optimization of a connectivity-based neurofeedback signal, we introduced and demonstrated a data-driven, process-based framework of designing a rtfMRI-nf treatment protocol with precise targeting and an optimally designed neurofeedback signal.

## 2. Materials and Methods

### 2.1. Data

Data of 268 participants including a mood and/or anxiety disorder group (MA; N = 223, 162 females, mean (SD) age = 36 (11) years, 147 participants were medicated) and a healthy control group (HC; N = 45, 23 females, mean (SD) age = 32 (11) years) from the Tulsa 1000 study (Victor et al., 2018) were used in the analysis. Of the note, the participants were selected from the first 500 subjects (exploratory dataset out of 1000 subjects study cohort) of Tulsa 10000 study. The diagnosis was based on an abbreviated version of the Mini International Neuropsychiatric Interview (MINI V.6.0) (Sheehan et al., 1998). MA group includes participants with either a mood disorder, an anxiety disorder, or both with MINI.

Rumination was evaluated with the Ruminative Response Styles (RRS) scale (Nolen-Hoeksema and Morrow, 1991). The total score of RRS as well as its sub-scores of depressive, brooding, and reflective rumination (Treynor et al., 2003) were used as a regressor for resting-state functional connectivity patterns in the MDMR analysis. Rumination is a multidimensional construct and its depressive and brooding components are considered maladaptive processes associated with disorder symptoms, while its reflective component could be an adaptive process (Watkins and Teasdale, 2004). Hence, the neuropathology associated with depressive and brooding rumination could be a target of intervention. We also used the depression and anxiety scales in the Patient Reported Outcome Measurement Information System (PROMIS) (Cella et al., 2010) to examine their effects on the connectivity pattern. The MDMR analysis was performed independently for each symptom scale.

Resting-state fMRI data were collected on a whole-body 3 Tesla MR750 MRI scanner (GE Healthcare, Milwaukee, WI) with an 8-channel receive-only head array coil at the Laureate Institute of Brain Research. Participants were instructed not to move and to relax and rest while looking at a cross on the screen during an 8-min resting-state scan. A single-shot gradient-recalled echo-planner imaging (EPI) sequence with sensitivity encoding (SENSE) was used with imaging parameters of TR/TE = 2000/27 ms, FA = 78°, FOV = 240 mm, 39 axial slices with 2.9-mm thickness without gap, matrix = 96×96, SENSE acceleration factor R = 2, sampling bandwidth = 250 kHz. The EPI images were reconstructed into a 128×128 matrix resulting in 1.875×1.875×2.9 mm^3^ voxel volume. For anatomical reference, T1-weighted MRI images with a magnetization-prepared rapid gradient-echo (MPRAGE) sequence with parameters of FOV = 240×190 mm, matrix = 256×256, 120 axial slices, slice thickness = 0.9 mm, 0.9375×0.9375×0.9 mm^3^ voxel volume, TR/TE = 5/2 ms, SENSE acceleration R = 2, flip angle = 8°, delay/inversion time TD/TI= 1400/725 ms, sampling bandwidth = 31.2 kHz, scan time = 5 min 40 s, were also acquired.

### 2.2. Connectome-wide association analysis

Preprocessing of functional images was performed with Analysis of Functional NeuroImages (AFNI) (http://afni.nimh.nih.gov/afni/). The initial five volumes were excluded from the analysis. The preprocessing included despiking, RETROICOR (Glover et al., 2000) and RVT (Birn et al., 2008) physiological noise corrections, slice-timing correction, motion corrections, nonlinear warping to the MNI template brain with resampling to 2mm^3^ voxels using the Advanced Normalization Tools (ANTs) (Avants et al., 2008) (http://stnava.github.io/ANTs/), smoothing with 6mm-FWHM kernel, and scaling to percent change relative to the mean signal in each voxel. General linear model (GLM) analysis was performed with regressors of 12 motion parameters (three rotations, three shifts, and their temporal derivatives), three principal components of ventricle signals, local white matter average signals (ANATICOR (Jo et al., 2010)), 4th-order Legendre polynomials for high-pass filtering, and censoring TRs with large head motion (> 0.25 mm frame-wise displacement). Voxel-wise residual signals of the GLM were used for the connectome-wide analysis.

Connectome-wide investigation for the association between functional connectivity (FC) and a symptom was performed with MDMR analysis (Anderson, 2001; Shehzad et al., 2014). We followed the procedure described in detail in Misaki et al. (2018a), and scripts for the analysis are available at GitHub (https://github.com/mamisaki/MDMR_fMRI). Briefly, the processed resting-state fMRI images were down-sampled to 4mm^3^ voxels, and then the voxels in gray matter regions were extracted. A connectivity map with Pearson correlations between signals was made from each voxel to all other voxels. The dependent variable of MDMR is a distance matrix of the connectivity maps between participants. The distance of the maps was calculated with Euclidean distance of Fisher’s z-transformed connectivity maps. The MDMR analysis evaluates the association between the distance matrix and the predictor variables with a linear model. The model includes a symptom score, group (MA/HC), and their interaction, as well as gender, medication status, age, and motion (mean frame-wise displacement) as covariates. The result of MDMR was represented with an *F* value that was the ratio of the variance explained by a certain regressor relative to the residual variance (Misaki et al., 2018a; Shehzad et al., 2014). A permutation test was performed to test the significance of the statistic, in which regressors of interest (symptom score, group, and their interaction) were orthogonalized with regard to nuisance regressors, and then the orthogonalized regressors of interest were randomly permuted (Winkler et al., 2014). Ten thousand random permutations were performed.

These procedures were repeated for individual seed voxels within the MPFC and PCC/Prec regions. The MPFC and PPC/Prec region masks were extracted from the DMN map obtained from 405 healthy participants’ resting-state fMRI data (Allen et al., 2014) provided at http://trendscenter.org/data/. One cluster in the MPFC and four clusters in the PCC and the precuneus in the DMN were used as the MPFC and PCC/Prec masks, respectively (Figure 1A). The MDMR analysis was performed for the MPFC and PCC/Prec regions, separately. The MDMR statistical map was thresholded with voxel-wise *p* < 0.005 and cluster-size corrected *p* < 0.05. Cluster-size corrected *p*-value was evaluated with the same permutation procedure as the voxel-wise test.

**Figure 1.**
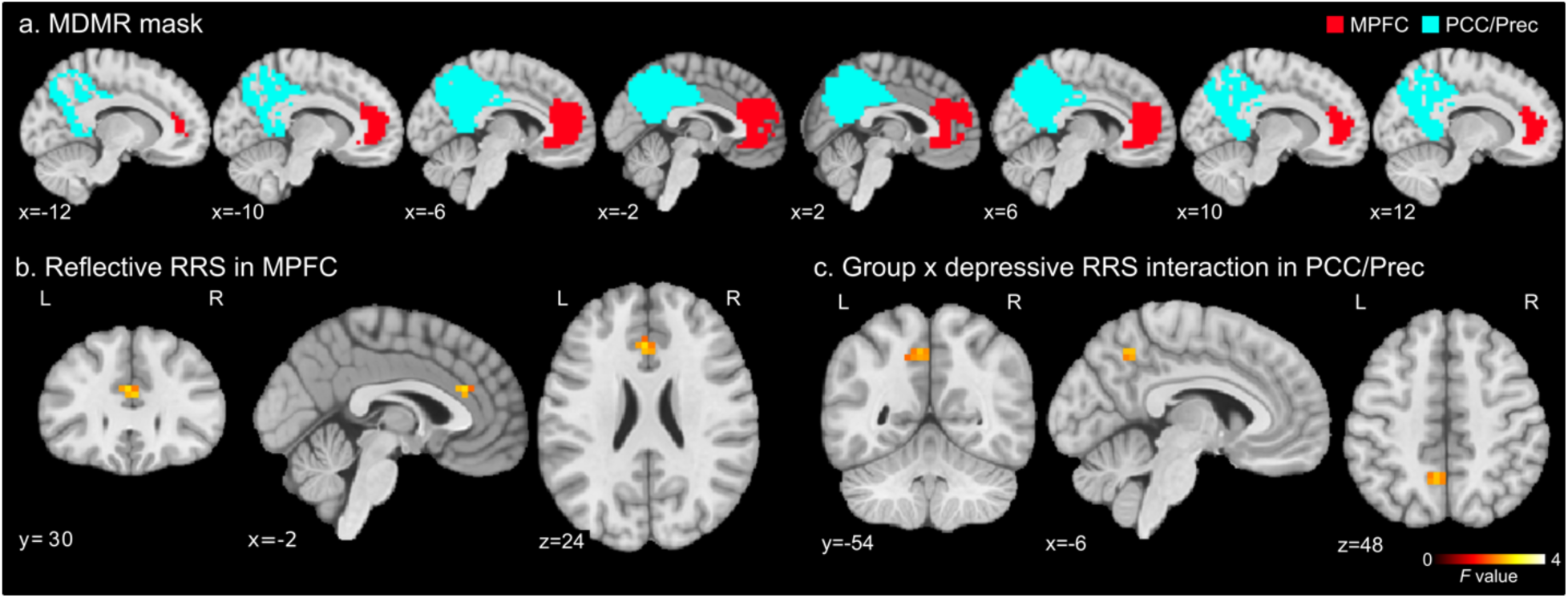
MDMR analysis mask and results. a) Masks of the medial prefrontal cortex (MPFC, red) and the posterior cingulate cortex/precuneus (PCC/Prec, cyan) regions, where MDMR analysis was performed. b) *F*-value map of the reflective RRS effect on the connectivity pattern in MDMR analysis in the discovered MPFC region. c), *F*-value map of the group by depressive RRS interaction effect on the connectivity pattern in MDMR analysis in the discovered PCC/Prec region. The maps were thresholded with voxel-wise *p* < 0.005 and cluster-size corrected *p* < 0.05.

Post-hoc seed-based connectivity analysis was performed for the discovered significant regions of the MDMR statistical map as the seed regions. The post-hoc analysis was done for the original resolution functional images. The seed region (6mm-radius sphere) was placed at a peak location of significant clusters in the MDMR statistical map. The average signal time-course of the seed area was used as a reference signal to calculate correlations with other voxels in the whole brain. Fisher’s *z*-transformation was applied to the correlation coefficient to make a connectivity map for each participant. Then, as a second-level group analysis, voxel-wise linear model analysis was applied to the connectivity maps with the same design matrix as the MDMR. The statistical map was thresholded with voxel-wise *p* < 0.001 and cluster-size correction of *p* < 0.05. The cluster-size threshold was evaluated with 3dClustSim in AFNI using a spatial autocorrelation function model (Cox et al., 2017).

Limiting the search area of MDMR helps increase the sensitivity of the analysis because the estimation of the null distribution derived from the permutation test used for family-wise error correction could be optimized for each region. This sensitivity improvement could be critical for the discovery because the sensitivity of MDMR analysis is lower than a seed-based analysis (Misaki et al., 2018b). We note that while the search for the seed voxels was limited in the masked regions, functional connectivity from the individual seed was evaluated for the whole brain voxels. Therefore, the analysis covered the whole brain connectivity that originated from the MPFC and PCC/Prec seed regions.

### 2.3. Simulation of online functional connectivity neurofeedback signal

To design an optimal neurofeedback signal for the identified FC associated with an RRS score, we performed a simulation to calculate an online real-time FC feedback signal. Here, two methods of online connectivity neurofeedback signal, sliding-window correlation (Gembris et al., 2000) and the two-point algorithm (Ramot et al., 2017), were evaluated. The sliding-window correlation is a *z*-transformed Pearson correlation between ROIs within a time window. Widths of a three-to ten-time points window were evaluated in the simulation. The window was moved at each TR to calculate the online feedback signal. The two-point algorithm uses the directionality of the signal change between the regions to evaluate their connectivity. The feedback signal is calculated as a binary value; e.g., when a participant is trained to increase the connectivity, positive feedback (+1) is given if the two regions have the same change direction (e.g., an increase or decrease); otherwise, no feedback (0) is given. The original introduction of the two-point method (Ramot et al., 2017) used a control ROI to cancel a signal change unspecific to the target connectivity. With the control ROI, positive feedback is given when the two target regions have the same change direction as well as that is different from the direction in the control region. Both versions of the two-point method, with and without the control ROI, were evaluated in the simulation.

Feedback signals of these online functional connectivity measures were calculated in a real-time fMRI processing simulation for the resting-state fMRI data used in the connectome-wide analysis. The simulation was performed on an advanced real-time fMRI data processing system implementing comprehensive online noise reduction processes (Misaki et al., 2015; Misaki and Bodurka, 2019). The system performed slice-timing correction, motion correction, spatial smoothing, signal scaling, and GLM with regressors of high-pass filtering, six motion parameters, mean white matter signal, mean ventricle signal, and RETROICOR (Glover et al., 2000) in real-time online processing. This system enabled us to obtain a cleaned online fMRI signal in real-time. The online FC was calculated for this online processed signal.

The optimality of online FC was evaluated with regard to three criteria, correlation with FC obtained from offline analysis, robustness to head motion, and timeliness of neurofeedback. Since the identified FC in the connectome-wide analysis had a significant association with rumination severity in the offline FC (correlation with whole time-course signals), an online FC that had a high correlation with the offline FC should be a better neurofeedback signal. Correlation between the offline FC and the average online FC neurofeedback signal time-course was calculated for this evaluation. Robustness to the head motion’s artifact was evaluated with the correlation between the time-course of the online FC neurofeedback signal and the time-course of the mean frame-wise displacement within the window of online connectivity calculation. For the two-point method, the window was the current and the previous time points. The timeliness of the online FC was determined by its dependence on the signal history. Dependence on history is large for methods with more time points; thus, a short-width sliding-window or the two-point method are preferred in this regard.

## 3. Results

### 3.1. Connectome-wide association analysis

Table 1 shows the demographic and symptom profile of the Tulsa 1000 participants used for this analysis. There was no significant difference in age between the groups. Symptom scales of rumination, depression, and anxiety were significantly higher for MA than HC. All the sub-scales of RRS were also significantly higher for MA than HC.

**Table 1.** Data demographic and symptom scales.

|  |  | HC | MA | *t*-test: MA-HC |
| --- | --- | --- | --- | --- |
|  |  | mean (SD) | mean (SD) |  |
| N |  | 45, 23 females | 223, 162 females |  |
| Age | | 32 (11) | 36 (11) | $t = 1.96, p = 0.054$ |
| RRS | Total | 29.8 (8.0) | 55.1 (11.5) | $t = 17.74, p < 0.001$ |
| | Depressive | 15.5 (4.7) | 31.2 (7.1) | $t = 18.41, p < 0.001$ |
| | Brooding | 6.8 (2.2) | 12.7 (3.4) | $t = 14.70, p < 0.001$ |
| | Reflective | 7.5 (2.7) | 11.2 (3.2) | $t = 8.09, p < 0.001$ |
| PROMIS | Depression | 43.7 (6.2) | 60.9 (7.7) | $t = 16.21, p < 0.001$ |
| PROMIS | Anxiety | 46.2 (7.8) | 62.8 (6.4) | $t = 13.45, p < 0.001$ |

HC: healthy controls; MA: mood and anxiety disorder; RRS: ruminative response style; PROMIS: Patient Reported Outcome Measurement Information System.

Figure 1a shows the masks for the MPFC and PCC/Prec regions, where the MDMR analysis was performed. A significant association between a symptom scale and the FC was found for reflective RRS and depressive RRS in the MDMR analysis. Specifically, a significant *F* value of the MDMR analysis for the effect of reflective RRS was found at the anterior cingulate cortex (ACC, x, y, z = −2, 30, 24 mm in MNI) in the MPFC (Fig. 1b). A significant effect of the group by depressive RRS interaction was found in the left precuneus (−6, −54, 48 mm in MNI) in the PCC/Prec (Fig. 1c). No other symptoms and their interaction with the group showed a significant effect on the connectivity pattern in the MDMR analysis. Post-hoc seed-based connectivity analysis was performed for the peak locations of the significant MDMR results.

Post-hoc analysis for the ACC seed connectivity revealed a significant effect of reflective RRS (Figure 2) at the bilateral fusiform and inferior temporal area, the bilateral middle frontal region, the left middle cingulate region, the left medial frontal region, the right precuneus, the right temporal pole, the right thalamus, and the right calcarine region. Peak coordinates of the significant clusters are shown in Table 2. The connectivity between the ACC and these regions were positively correlated with reflective RRS.

**Figure 2.**
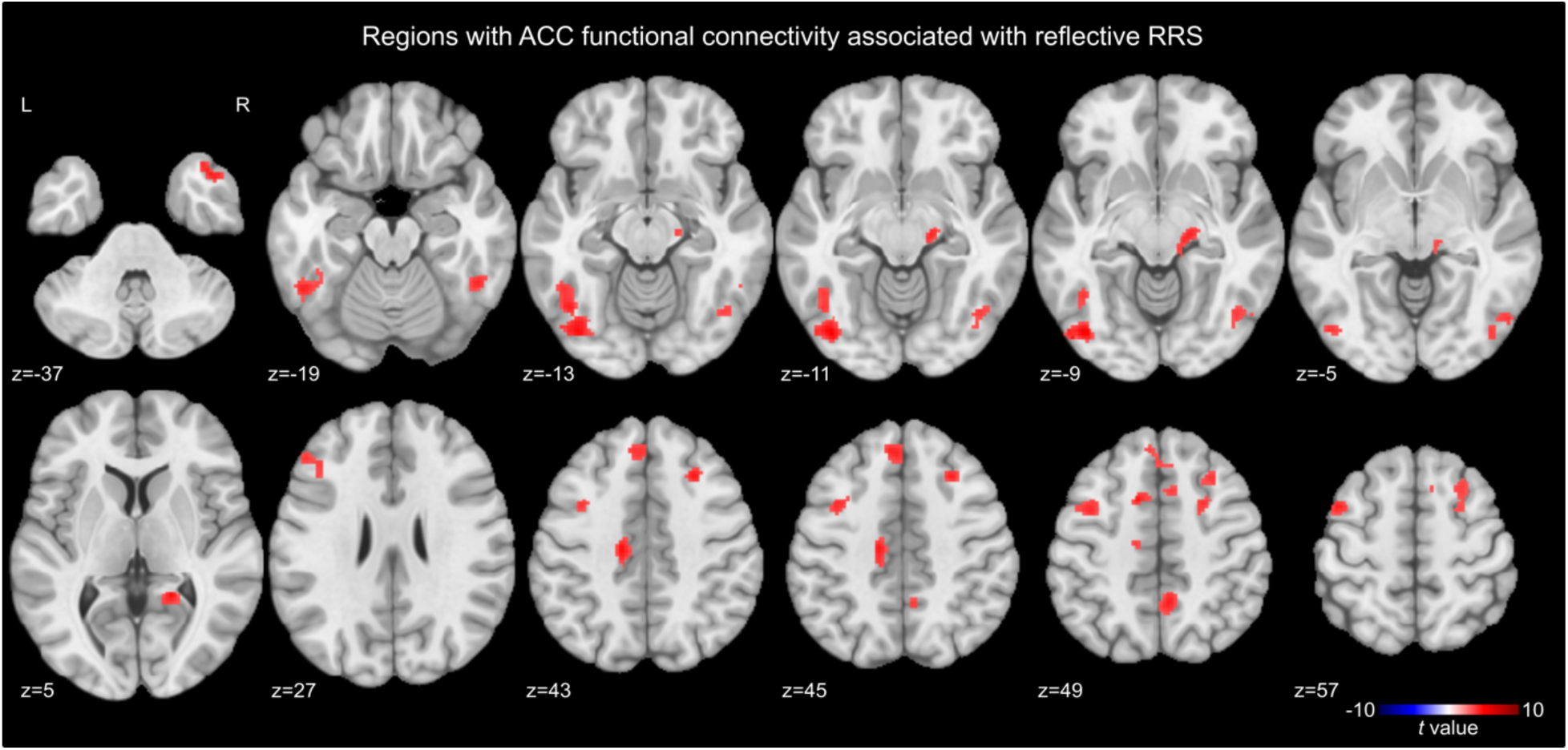
Regions with anterior cingulate cortex (ACC) connectivity significantly associated with reflective RRS. The map was thresholded with voxel-wise *p* < 0.001 and cluster-size corrected *p* < 0.05.

**Table 2.** Peak coordinates (MNI) of clusters with the significant effect of reflective RRS on the anterior cingulate cortex (x, y, z = −2, 30, 24 mm) functional connectivity.

| Region | x | y | z | t-value |
| --- | --- | --- | --- | --- |
| Left fusiform | -43 | -77 | -9 | 5.072 |
| Right middle frontal | 27 | 23 | 43 | 4.578 |
| Left middle Frontal | -45 | 5 | 57 | 4.316 |
| Left middle cingulate | -15 | -19 | 43 | 4.682 |
| Left medial Frontal | -13 | 9 | 49 | 4.758 |
| Right inferior temporal | 53 | -69 | -5 | 3.847 |
| Left middle frontal | -49 | 33 | 27 | 3.917 |
| Left medial frontal | -5 | 37 | 45 | 4.336 |
| Right precuneus | 5 | -51 | 49 | 4.399 |
| Right temporal pole | 43 | 15 | -37 | 4.503 |
| Right fusiform | 49 | -49 | -19 | 4.237 |
| Right thalamus | 17 | -21 | -11 | 4.268 |
| Right calcarine | 19 | -47 | 5 | 4.697 |

Post-hoc analysis for the effect of depressive RRS on the precuneus connectivity was performed for each of the HC and MA groups to resolve the significant interaction effect on the precuneus connectivity. Figures 3a and 3b show the regions with the precuneus connectivity significantly associated with depressive RRS for HC and MA, respectively. The HC group had a significant association between depressive RRS and precuneus connectivity at the right precentral and paracentral regions, the right supplementary motor area (SMA), the bilateral insula, and the left inferior occipital region (Fig. 3a). Connectivity between the precuneus and these regions was negatively correlated with depressive RRS in HC. The MA group had a significant association between depressive RRS and the precuneus connectivity at the right temporoparietal junction (TPJ) area and the left intraparietal sulcus (IPS) region (Fig. 3b). Connectivity between the precuneus and these regions was positively correlated with depressive RRS in MA. Peak coordinates of the significant clusters are shown in Table 3.

**Figure 3.**
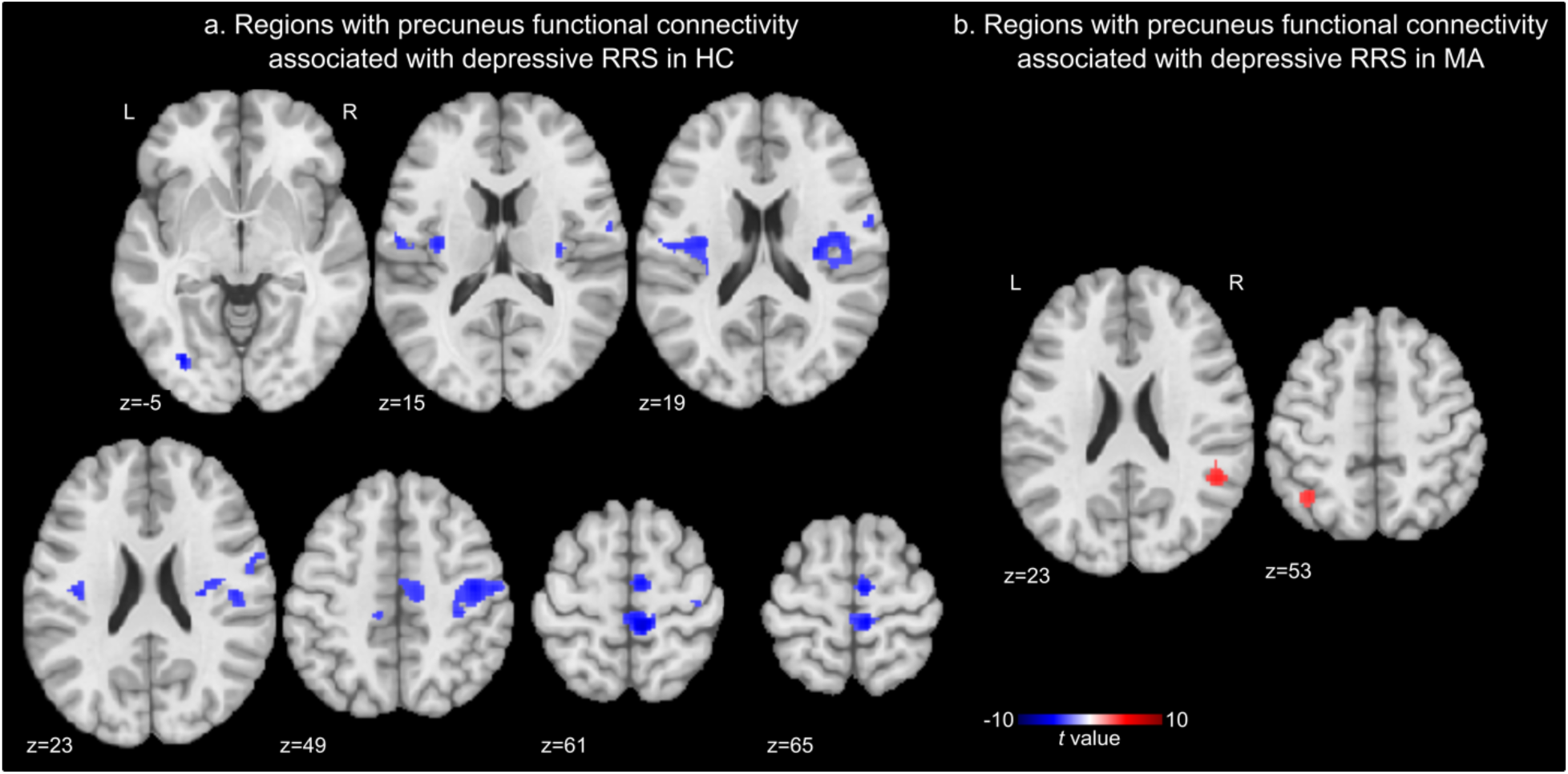
Regions with precuneus connectivity significantly associated with depressive RRS in healthy control (HC) (a) and mood and anxiety disorder (MA) (b) group. The maps were thresholded with voxel-wise *p* < 0.001 and cluster-size corrected *p* < 0.05.

**Table 3.**
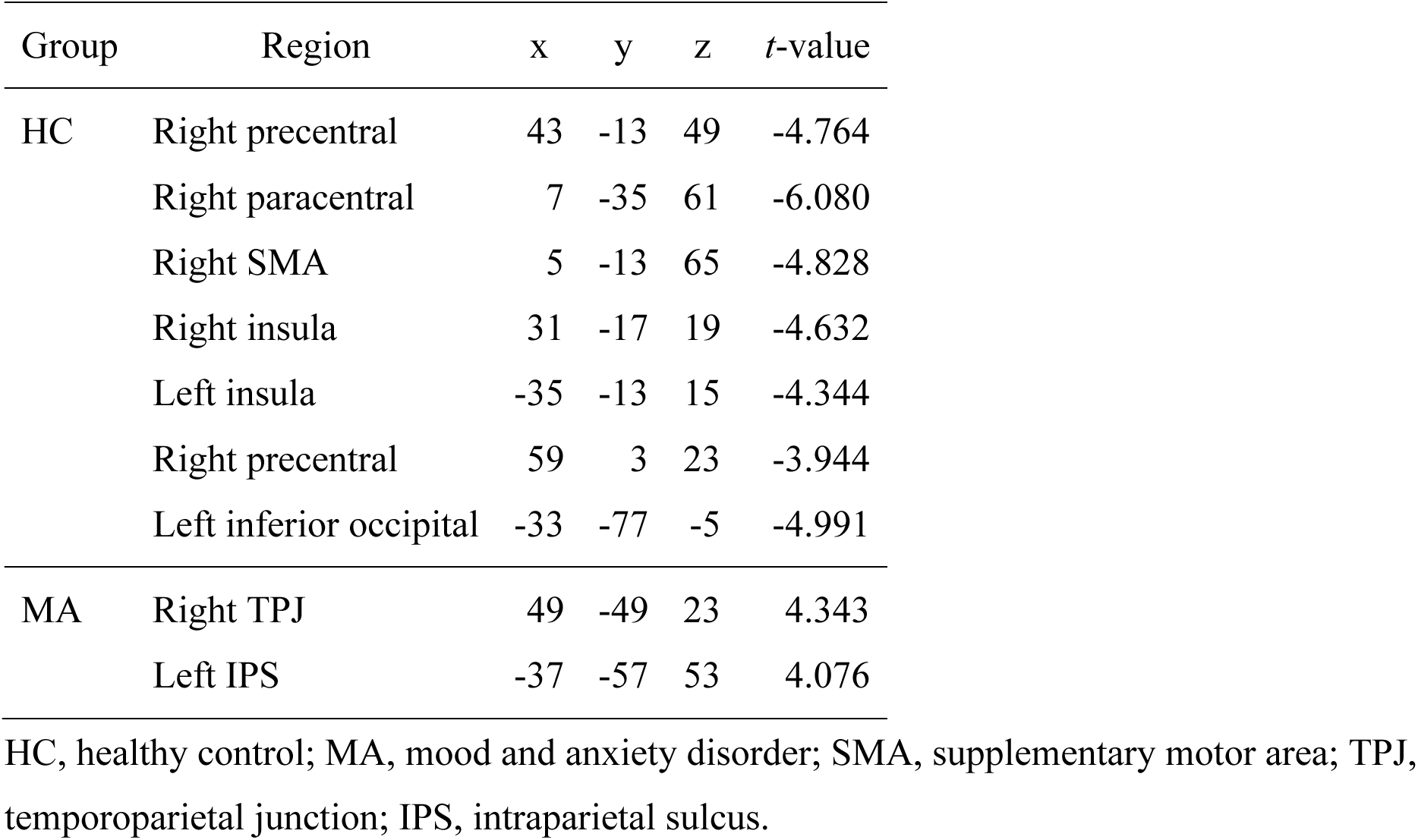
Peak coordinates (MNI) of clusters with a significant effect of depressive RRS on the precuneus functional connectivity for healthy and mood/anxiety disorder groups.

These results indicated that pathological rumination in MA was associated with the precuneus connectivity with the right TPJ and the left IPS regions. We further examined the association with other RRS subscales as well as the effect of gender and medication status on these connectivities. Figure 4 shows associations between the precuneus connectivity and RRS subscales for the right TPJ and the left IPS regions. Connectivity (*z*-transformed Pearson correlation) was calculated between the mean signals of a 6-mm-radius sphere ROI centered at the peak locations. The right TPJ connectivity with the precuneus had significant associations with all RRS subscales in the MA group. Left IPS connectivity with the precuneus had a significant association only with depressive RRS in MA. No significant effect of gender was found on either connectivity when the interaction of gender by depressive RRS was added in the analysis. When the interaction of medication status by depressive RRS was added in the analysis, a significant interaction effect of medication by depressive RRS was observed for the right TPJ connectivity (*F* = 4.663, *p* = 0.031). This effect was driven by the larger association in unmedicated (*t* = 4.189, *p* < 0.001) than medicated participants (*t* = 1.986, *p* = 0.048). No significant interaction effect of the medication on depressive RRS was found for the left IPS connectivity (*F* = 0.382, *p* = 0.537). Considering that all RRS subscales were significantly higher for MA than HC (Tab. 1), these results suggest that the precuneus connectivity with the right TPJ could be more strongly associated with the severity of rumination symptoms in MA disorder than with the left IPS.

**Figure 4.**
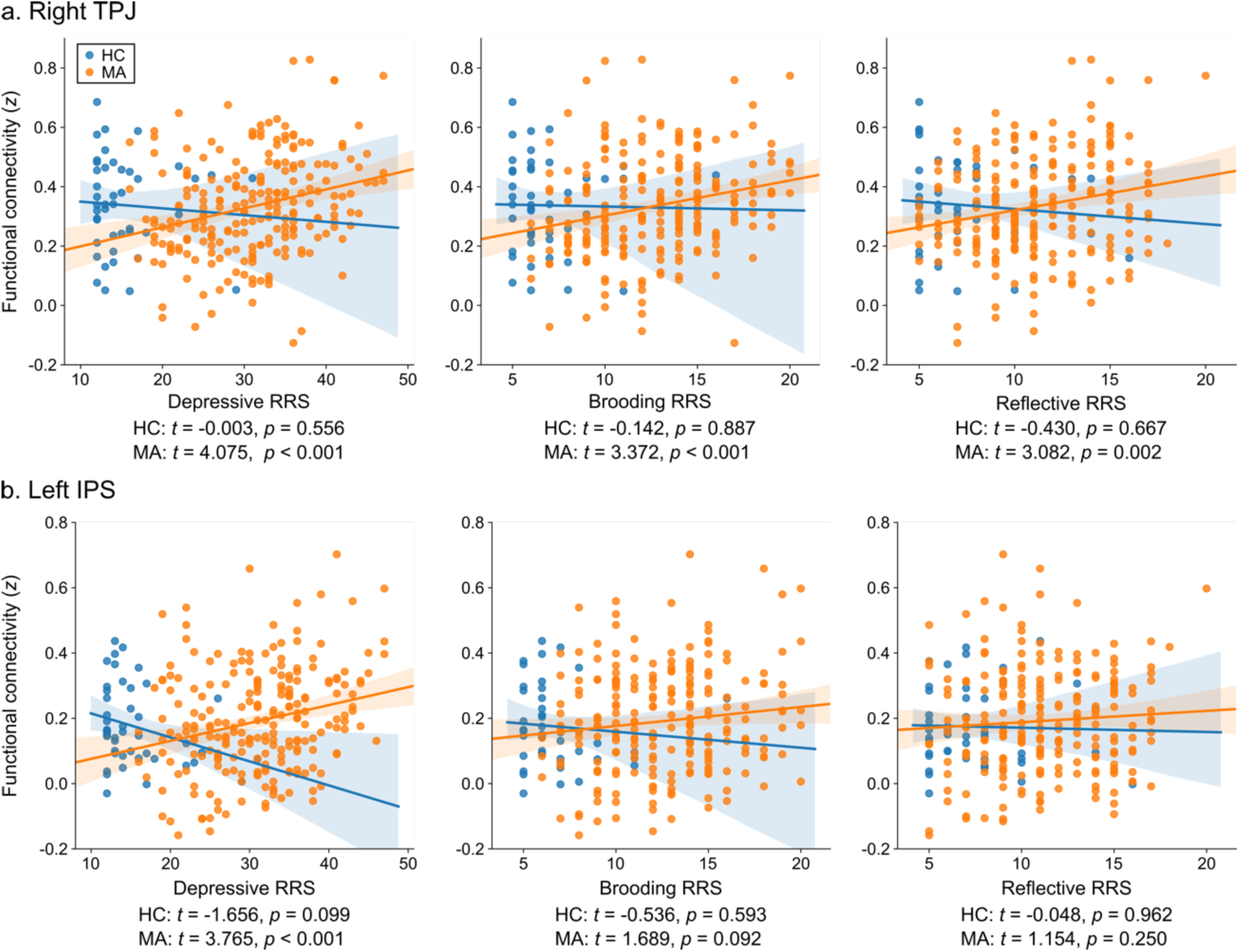
Associations between the precuneus connectivity and RRS subscales in the right temporoparietal junction (TPJ, a) and the left intraparietal sulcus (IPS, b) regions for the healthy control (HC) and mood and anxiety disorder (MA) groups. Each point indicates an individual participant. Fitted lines and their 95% confidence intervals for HC and MA are also shown. *t* and *p* values indicate the significance of the linear association between RRS and connectivity for each group.

### 3.2. Simulation of online functional connectivity neurofeedback signal

The FC between the precuneus and the right TPJ was determined as a promising rtfMRI-nf target to relieve the rumination symptoms. We, therefore, performed a simulation analysis of online FC neurofeedback signal for this connectivity. Control ROI for the two-point method was placed at the right precentral region (6mm-radius sphere centered at x, y, z = 31, −29, 69 mm in MNI), where the correlation between motion and its connectivity with the precuneus was highest in the cortex. Thus, controlling the signal change in this ROI could remove the effect of motion on the FC neurofeedback signal.

Figure 5a shows the correlations between the offline and online FCs with the two-point and sliding-window methods in a real-time fMRI processing simulation. Including the control point in the two-point method decreased the correlation, and the more time points the online calculation included, the higher the correlation with the offline measure was observed. Figure 5b shows the distribution of correlation between head motion and online connectivity measures across participants. The plot indicates that the more time points the online calculation included, the more participants had a high absolute correlation with motion.

**Figure 5.**
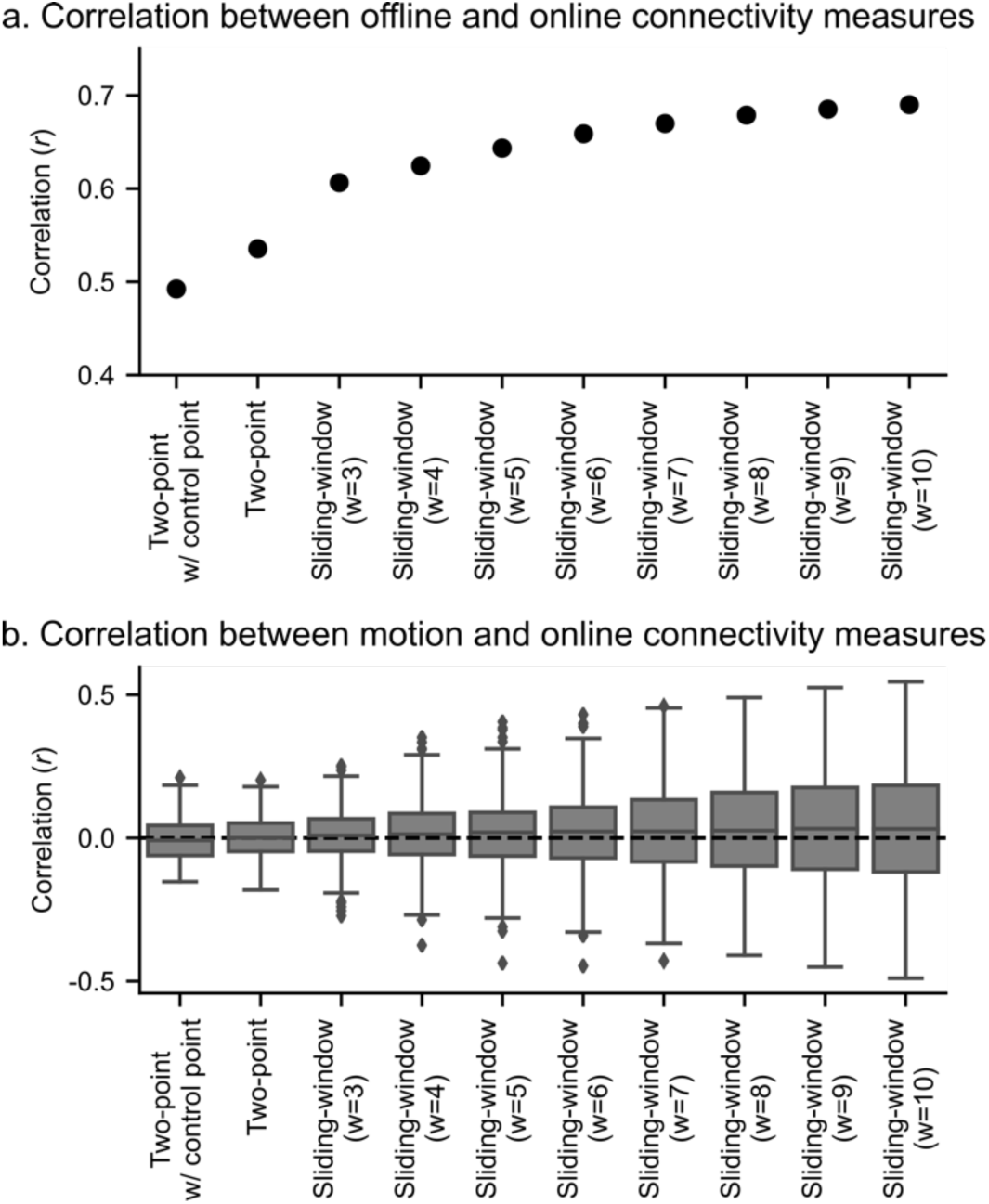
The results of the real-time fMRI processing simulation. a, Correlation between the offline and online connectivity measures for the two-point and sliding-window methods with different window width (w). b, Distributions of correlations between head motion and online connectivity measures across participants. The box shows the range of the 1st to 3rd quartile values (inter-quartile range, IQR) and the extending lines indicate the whole range of values except outliers (values larger or less than 1.5 x IQR from the edge of the box), which are indicated with dots. The line in the box shows the median value.

## 4. Discussion

We performed a data-driven process-based search for a rtfMRI-nf target of functional connectivity associated with rumination symptoms using resting-state fMRI data on the cohort of mood and anxiety disorder individuals and healthy controls. The connectome-wide association analysis revealed that the connectivity between the precuneus (−6, −54, 48 mm in MNI) and the right TPJ (RTPJ; 49, −49, 23 mm in MNI) was significantly associated with depressive RRS as well as brooding and reflective RRS in MA, and this association was greater for unmedicated than medicated participants. The simulation analysis of the online FC neurofeedback signal for this connectivity indicated that while the methods with more time points had a high correlation with offline FC, these also had a high risk of contamination by motion. In addition, a method with more time points is also not favored in regard to the timeliness of feedback signal.

While depressive and brooding components, rather than reflective component of RRS have been associated with the pathological effect of rumination (Watkins and Teasdale, 2004), the current data indicated that MA participants had significantly high reflective RRS as well as depressive and brooding RRS. High RRS in both reflective and brooding components was also observed in another study (Satyshur et al., 2018). This observation was consistent with the study indicating that brooding and reflective rumination were not separate factors in depressed patients because these components could exacerbate each other in depression (Whitmer and Gotlib, 2011). The significant effect of depressive RRS was found in the FC between the precuneus and RTPJ and between the precuneus and the left intraparietal sulcus (LIPS) region. The precuneus– RTPJ FC was significantly associated with all components of RRS, while the precuneus–LIPS FC was significantly associated only with depressive RRS. Considering that all components of RRS were significantly higher for MA than HC, and each component of RRS could exacerbate each other in a pathological state, the precuneus–RTPJ functional connectivity constitutes a very promising intervention target for FC-based rtfMRI-nf to treat pathological rumination.

The TPJ region, especially in the right hemisphere, has been implicated in attentional and social functions, including the theory of mind and self-other judgment (Eddy, 2016). Specifically, its anterior part is associated with externally-oriented, stimulus-driven attention with high connectivity to attentional selection regions, while its posterior part is associated with internally-oriented, stimulus-independent process with high connectivity to regions for social cognitions (Bzdok et al., 2013; Mars et al., 2012; Uddin et al., 2010). The RTPJ region in the present result was included in the posterior part of TPJ in either parcellation of anatomical connectivity of diffusion tensor imaging (Mars et al., 2012), the task-related meta-analytic connectivity, or resting-state functional connectivity (Bzdok et al., 2013). Meta-analysis studies also found that brain activations for social cognitive tasks overlapped with the DMN regions in the dorsomedial PFC, the precuneus, and the TPJ areas (Schilbach et al., 2012; Spreng et al., 2009; Van Overwalle, 2009). These regions coactivated in autobiographical memory and theory of mind tasks (Spreng et al., 2009). These suggest that high connectivity between the precuneus and RTPJ could be associated with thinking of autobiographical events in a social context that might be associated with exacerbation of negative judgment in rumination. Coactivation of the precuneus and the TPJ was also reported for the contrast between other- and self-agency conditions, with higher activation at the other-agency condition (Farrer and Frith, 2002; Murray et al., 2015). Hence, critical thinking of autobiographical things with others’ viewpoints, which could amplify rumination severity, might be associated with a high precuneus–RTPJ connectivity.

This connectivity, however, has not been identified in studies investigating FCs associated with self-referential processing and rumination symptoms. Actually, several studies suggest that MPFC was involved in self-referential processing and rumination. The MPFC implication in self-referential thinking has been indicated in a direct examination of a self-referential thinking task, which showed that MPFC activity was high in self-referential thinking relative to a general thinking condition (Nejad et al., 2019). Interestingly, while RRS was positively correlated with this activity in healthy participants, it was negatively correlated with RRS in remitted MDD participants. This decreased MPFC activity associated with high RRS in participants with a high risk of MDD suggests that the self-recognition process decreased in the ruminative state. If the present result of increased precuneus–RTPJ connectivity is linked to thinking with others’ viewpoints (Farrer and Frith, 2002; Murray et al., 2015), the decrease of MPFC activity associated with a decreased self-recognition process might be consistent with the current result. Thus, the precuneus-RTPJ FC might not be implicated in the self-referential processing itself but could be related to the process of exacerbating and maintaining pathological rumination through thinking autobiographical events with critical others’ perspective.

Alterations in FC associated with rumination have also been indicated for MPFC seed connectivity. Increased resting-state FC between the MPFC seed and the PCC was correlated with reflective rumination for female participants with major depressive disorder (MDD) (Satyshur et al., 2018). This result is partly consistent with the present result that found the association between reflective RRS and the MPFC connectivity (Fig. 2, Tab. 2). Resting-state FC between the dorsal MPFC seed and the temporal pole was positively correlated with depressive rumination for first-episode treatment-naïve young adults with MDD (Zhu et al., 2017). Resting-state FC between the MPFC seed and the left inferior parietal lobule was also positively correlated with the severity of negative self-focused thought (Philippi et al., 2018). Positive correlation with negative self-focused thought was also seen for pregenual ACC seed connectivity with the dorsolateral PFC, the precuneus, the inferior parietal cortex, and the paracentral lobule extending to SMA (Philippi et al., 2018). Resting-state FC for the subgenual ACC with the right middle and inferior frontal gyrus was negatively correlated with RRS (Connolly et al., 2013). These results suggest that MPFC is implicated in self-referential processing and the rumination process, while the associated FC was not consistent.

We should note that the results of these FC association with rumination studies cannot be compared to the current result because those used *a priori* defined seed ROI, and the analysis for the RRS association was performed post hoc for the FC with a significant difference between the depressed and healthy groups. Thus, even though the current effect has not been found in those studies, they do not contradict the current result. Also, while the present analysis did not find associations between depressive rumination and FC in the MPFC, this could be explained by the low sensitivity of MDMR analysis relative to a seed-based connectivity analysis (Misaki et al., 2018b). Since the MDMR analysis could find a significant association only with a large effect, the found association between the precuneus-RTPJ connectivity and the RRS should be considered a robust result. In addition, the aim of this analysis was to discover the rtfMRI-nf intervention target, not to describe the connectivity affected by the rumination comprehensively. Thus, finding the FC with a robust association with rumination should be suitable for the present purpose.

Additionally, we performed simulation analysis to find the optimal online functional connectivity neurofeedback signal for the precuneus-RTPJ connectivity. The real-time fMRI processing simulation indicated a trade-off between the correlation with offline FC and the risk of motion contamination. The higher correlation with the offline FC for the methods with more time points was not surprising because the offline FC includes all time points for its calculation. The higher correlation with motion was because the methods with more time points had prolonged dependence on signal history, which could spread the effect of a time point with a significant head motion to many points of the neurofeedback signal. Dependence of long signal history is also unfavored regarding the timeliness of the feedback signal. Because a motion artifact could be critical as a risk of implicit learning of artifact effect in rtfMRI-nf training (Zhang et al., 2011), and the online rtfMRI-nf signal should minimize the delay as the fMRI signal already includes the hemodynamic response delay, a feedback signal with fewer time points should be preferred. While the correlation with offline FC was lower for the methods with fewer time points, the observed correlation, higher than 0.5 (Fig 5a), could be high enough to train the participants to regulate the target connectivity. As a matter of fact, many rtfMRI-nf studies demonstrated successful self-regulation training even without a comprehensive real-time noise reduction process (Heunis et al., 2018), and the correlation between the online- and offline-calculated signals was not high when the real-time noise reduction process was not comparable to the offline one (Misaki and Bodurka, 2019). When we performed a simulation with only the motion correction in real-time processing – which is a conventional rtfMRI process used in many studies – the correlation between the online and offline FC was less than 0.5 even for the highest online FC neurofeedback signal (10-TR sliding-window). This indicates that an online neurofeedback signal with a correlation as high as 0.5 with offline-evaluated FC could be enough to train a participant to regulate the target brain activity. Taken together, we consider the two-point method or a sliding-window correlation with short window width as favored to an online FC neurofeedback signal.

For the two-point method, using the control ROI did not help to reduce the motion effect compared to the two-point method without the control ROI (Fig. 5b). This result could be attributed to the fact that the current real-time fMRI processing simulation included a comprehensive noise reduction process; thus, the additional benefit of controlling motion was minimal. Using the control ROI also decreased the correlation with offline FC, which could be due to a reduced positive feedback frequency with restriction by the control ROI.

The connectome-wide investigation indicated that the precuneus-RTPJ connectivity was positively correlated with rumination severity. Hence, the rtfMRI-nf training to treat rumination symptoms should train a participant to decrease this connectivity. At the rtfMRI-nf training to decrease FC, the two-point method is more convenient than the sliding-window correlation because the sliding-window correlation requires a baseline level of connectivity to present feedback for signal reduction. Defining baseline connectivity is not a trivial task, though - it will need a personalized approach. In contrast, the two-point method does not need a baseline setting because the feedback signal is a binary value. We can give positive feedback when the two regions have different change directions, and give no feedback when they are the same. In light of these considerations, we suggest that the two-point method without control ROI is the most convenient, robust to motion, and timely online FC neurofeedback method to train a participant to decrease FC as far as we use the comprehensive real-time fMRI noise reduction process.

Several limitations of this study should be acknowledged. Although the association between RRS and the precuneus-RTPJ connectivity was seen specifically for the MA group, this specificity could be due to the unbalanced number of participants. The present result might be biased to the MA population, and may not be generalized to rumination for the preclinical population. With the limitation of the sensitivity of MDMR, the analysis could detect only the association with a large effect. MDMR analysis is insensitive to a change in a small region because the analysis depends on the between-subject distance matrix, which summarizes the difference between a whole-brain connectivity maps into one distance measure (Misaki et al., 2018a, b). Permutation test used in the analysis also limits the sensitivity (Misaki et al., 2019). As the bias-variance trade-off in model complexity suggested (Bishop, 2007), null distribution in the permutation test could have large variance with a large multivariate model fitted to a limited number of samples, which makes it hard to find a significant effect. Therefore, the absence of a significant effect in other symptoms does not prove the absence of their effect on resting-state FC. Limiting the MDMR search within the MPFC and PCC/Prec areas could also limit the findings. While limiting the search region has methodological merit in increasing the analysis sensitivity, and these areas are the most credible regions for searching for association with rumination, there might be FCs that do not stem from these areas but have a strong association with rumination symptoms. Nevertheless, the identified association between RRS and the precuneus-RTPJ connectivity was significant, and this connectivity is a promising target of rtfMRI-nf that can possibly treat rumination symptoms. We should remember that the present result did not describe a comprehensive abnormality of resting-state FC associated with rumination but rather discovered the credible target of rtfMRI-nf intervention to treat rumination symptoms.

## 5. Conclusion

The data-driven process-based approach discovered the functional connectivity locus in the precuneus associated with rumination severity. We showed that the precuneus-RTPJ connectivity is a promising target of rtfMRI-nf intervention to treat rumination symptoms. The simulation analysis of the online FC neurofeedback signal suggested that the two-point method without control ROI was robust to motion, less dependent on the signal history, and convenient for the training to decrease FC. In future studies, we will examine the utility of rtfMRI-nf training to reduce the precuneus-RTPJ connectivity with the two-point method for alleviating pathological rumination.

The present study offers a discovery framework for the precise location of functional connectivity targets for rtfMRI-nf intervention. This framework could identify the target with high spatial specificity and is applicable to a wide range of symptom dimensions. In the future, the current approach could help rtfMRI-nf become fully-fledged as a clinical treatment, with a direct application of the neuroimaging result to clinical interventions focused on improving psychiatric symptoms and modifying the trajectory of psychiatric disorders.

## Acknowledgement

This work has been supported in part by The William K. Warren Foundation and the National Institute of General Medical Sciences Center Grant Award Number 1P20GM121312. The content is solely the responsibility of the authors and does not necessarily represent the official views of the National Institutes of Health. The Tulsa 1000 Investigators include the following contributors: Robin Aupperle, Ph.D., Jerzy Bodurka, Ph.D., Justin Feinstein, Ph.D., Sahib S. Khalsa, M.D., Ph.D., Rayus Kuplicki, Ph.D., Martin P. Paulus, M.D., Jonathan Savitz, Ph.D., Jennifer Stewart, Ph.D., Teresa A. Victor, Ph.D.

